# Pax3 repairs a neural circuit through a program of directed axon outgrowth

**DOI:** 10.1101/2021.02.25.432898

**Authors:** J. Sebastian Jara, Hasan X. Avci, Ioanna Kouremenou, Mohamed Doulazmi, Joelle Bakouche, Caroline Dubacq, Catherine Goyenvalle, Jean Mariani, Ann M. Lohof, Rachel M. Sherrard

## Abstract

Repairing damaged or dysfunctional human brain circuits remains an ongoing challenge for biomedical science. While surviving neuronal networks can be reorganised after lesion, for example by neurotrophins, these new connections are disorganised and rarely produce clinical improvement. Here we investigate how to promote axonal growth while retaining correct cellular targeting. We show that, in response to brain-derived neurotrophic factor (BDNF) in target-tissue, potential reinnervating neurons upregulate Pax3. Pax3 in turn increases polysialic acid-neural cell adhesion molecule (PSA-NCAM) on their axon terminals, facilitating their outgrowth and pathfinding, and resulting in correctly-targeted neural circuit repair in the mature nervous system. This is a novel role for Pax3, which we confirmed by showing its expression in afferent neurons is essential for spontaneous and BDNF-induced reinnervation in the developing and mature brains, respectively. Together these results suggest that Pax3 contributes to a repair program, in which axon growth is promoted *and* direction signaling maintained. These data advance our understanding for accurately rebuilding neural circuits: restricting growth-promotion to potential afferent neurons, as opposed to stimulating the whole circuit, allows axon growth without impairing its guidance.

## INTRODUCTION

Repairing neuronal circuit damage or dysfunction remains a major challenge in biomedical science, because the poor growth properties of adult neurons and inhibitory extracellular molecules prevent effective axon outgrowth ^**1**,**2**^. Moreover, neurons are lost during trauma or neurodegenerative disease, so re-growing an identical circuit is not possible, and functional recovery requires the structural reorganisation of remaining undamaged axons ^**3**,**4**^. Such post-lesion reorganisation of residual circuits is a phenomenon of the developing brain ^**5–7**^. Unfortunately, treatment strategies to recreate developmental processes, by increasing a neuron’s growth capacity ^**8**,**9**^, or the growth-permissiveness of the cellular environment ^**10**,**11**^, generally produce inaccurate connections that do not restore circuit function and may even have deleterious effects ^**12**^. It is therefore imperative to identify molecules that can promote axon growth while also allowing appropriate cellular targeting and synaptogenesis as would occur during development.

The neurotrophin family of growth factors has fundamental roles in neural development and consequently has been widely studied in neural circuit repair. While neurotrophins may improve neuronal survival and axon sprouting, there is little clinical benefit ^**13**,**14**^. It is only when neurotrophins, in particular BDNF (brain-derived neurotrophic factor), are examined in their developmental roles as target derived factors ^**15**,**16**^, that axon collateral growth effectively reorganises spared connections to improve function. The effect of BDNF is highly dose-dependent; repeated high doses induce excess terminal sprouts that cause pathological pain and spasticity ^**12**,**17**^, whereas more physiological concentrations have fewer adverse effects ^**12**,**18**^, but are also less effective. To promote beneficial postlesion plasticity and avoid unwanted effects, identifying repair mechanisms downstream of BDNF may allow a more focussed action without excessive sprouting, and thus fewer inaccurate neural connections.

In this study we examined molecular pathways downstream from BDNF using an experimental model in which the biological effect can be readily quantified, and in which we can separate treatments to act *either* on axonal outgrowth (presynaptic) *or* by target attraction (postsynaptic). This is post-lesion repair of the olivocerebellar path, in which reinnervating olivary axons (climbing fibres) are topographically organised and correctly targeted at cellular and subcellular levels ^**7**,**19–21**^. In our *ex vivo* model of this path ^**19**,**22**^, we show that BDNF treatment to the target cerebellum increases expression of the transcription factor Pax3 and the growth-permissive molecule, polysialic acid-neural cell adhesion molecule (PSA-NCAM), *only* in the (afferent) inferior olive. Although the molecular links between cerebellar BDNF and changes in olivary gene expression are not known, olivary expression of either Pax3 or PSA-NCAM is required for reinnervation to take place, and upregulation of either molecule induces correctly targeted climbing fibre outgrowth. Although Pax3 is strongly expressed in the embryonic hindbrain, where it regulates regional developmental genes ^**23**^, it has not previously been linked to axonal outgrowth in the mammalian brain. These data demonstrate that molecules downstream of BDNF, acting on a subset of mechanisms (e.g. axonal outgrowth *or* target attraction), may provide a more-focussed and therefore more effective approach to produce circuit repair. This information may facilitate the development of new treatment strategies for human neurological disease.

## RESULTS

We used denervated co-cultured hindbrain explants (Figure S1) to elucidate mechanisms downstream of BDNF that underlie appropriately-targeted neural circuit repair. In this model, target-derived BDNF induces intact host climbing fibres to grow collaterals into a graft hemicerebellum and reinnervate Purkinje cells ^**19**,**22**^.

### BDNF injection into the graft hemicerebellum increases terminal sprouting from intact (host) climbing fibres

First we confirmed that olivocerebellar reinnervation *ex vivo* follows the same maturational changes as *in vivo*: spontaneous reinnervation during development (lesion at 9 DIV, ∼P3) and BDNF-induced reinnervation in the maturing system (lesion at 21 DIV, ∼ P15; ^**7**,**20**^). Cultured cerebellar plates (graft) were removed from their brainstem (denervation) and apposed to the cerebellar plates of an intact explant (host) for reinnervation (Fig S1A; ^**19**,**22**^) and those lesioned at 21 DIV were treated with BDNF or vehicle. After 10 days co-culture, we observed VGLUT2-labelled terminals around Purkinje cell somata and dendrites (Fig S1B) in hemicerebella lesioned at 9DIV, or 21DIV treated with BDNF, consistent with functional climbing fibre reinnervation ^**22**^. There was no such labelling in vehicle-treated 21DIV lesion hemicerebella. Reinnervation density was greater after lesion at 9 DIV than at 21 DIV, and at both ages reinnervation decreased with increasing laterality into the hemicerebellum (Fig S1C), as demonstrated *in vivo* ^**7**,**15**^. This suggests that cerebellar factors, such as BDNF, act on the olivary climbing fibre afferents to induce their outgrowth.

We therefore studied the initial effects of BDNF on host climbing fibre terminals that might explain their subsequent growth and reinnervation of graft Purkinje cells. We injected the inferior olive with lentivirus expressing GFP (LV-GFP) to visualize climbing fibre arbors in the host hemicerebellum (Fig 1A) and compared their structure in intact explants (control) and explants co-cultured at 21 DIV treated with exogenous BDNF (or vehicle). Host climbing fibre arbors were modified 24h after co-culture. Compared to vehicle-treated or control (no apposed graft) explants (Fig 1B), BDNF-treated arbors had many thin long horizontal branches. In particular, BDNF treatment resulted in more climbing fibres with horizontal branches greater than 40 µm long (Fig 1B&C).

**Figure 1.**
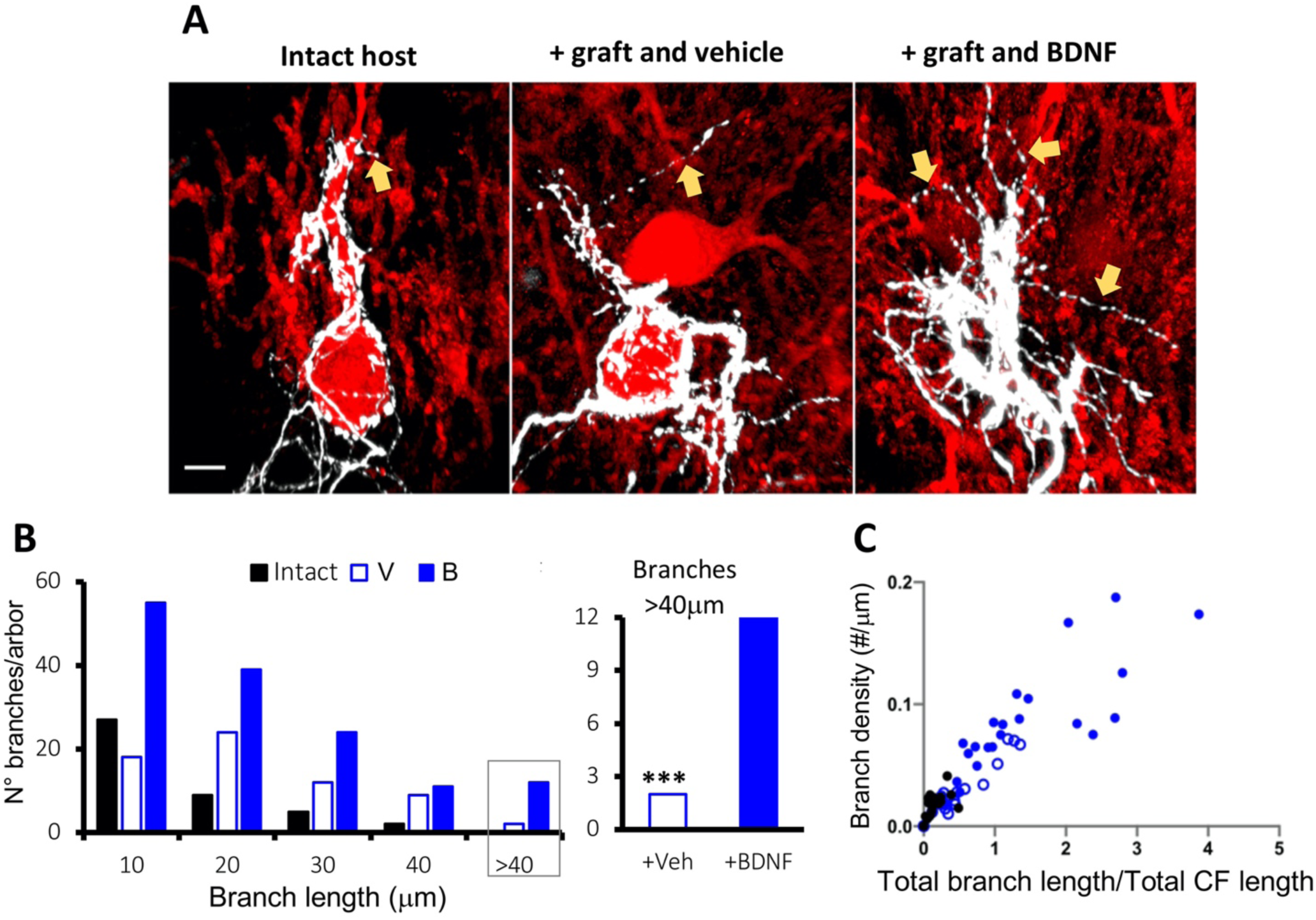
BDNF administration rapidly increases collateral sprouting from host CFs. **A:** In the intact host cerebellar plate (left panel), a few horizontal branches (gold arrow) project from GFP-expressing olivary climbing fibres innervating Purkinje cells. After lesion and coculture (middle panel) labelled climbing fibres of the *intact host* hemicerebellum have longer thin horizontal branches (gold arrow), and in the presence of BDNF (right panel) these horizontal branches of intact climbing fibres are also more numerous (gold arrows). **B:** Quantification of horizontal climbing fibre branches in the intact host hemicerebellum 24 hours after lesion and coculture. Horizontal branches are more numerous (left panel; ANOVA F2,57 = 12.7, p<0.0001) and longer (ANOVA F2,52 = 19.93, p<0.0001) in cocultures treated with BDNF (B). The number of horizontal branches longer than 40 µm was significantly higher after BDNF application (B) than vehicle-treatment (V) or intact (non-lesioned; intact) controls (χ ^2^ = 29.32, 2df, p<0.0001). **C:** Climbing fibre horizontal branch morphology is illustrated by plotting branch density vs. total branch length/total axonal length ^**52**^. In comparison with normal climbing fibre horizontal branches in the intact cerebellar plate, BDNF increases the density of longer branches Between group post-hoc comparisons BDNF vs. vehicle: ***, p<0.001. Bar = 20 µm.

To test whether this sprouting was due to diffusion of BDNF into the host hemicerebellum to affect these intact climbing fibre terminals, we used c-fos immunolabelling 1½h after treatment to identify BDNF-stimulated cells. C-fos labelling was only detectable in the BDNF-treated *graft* cerebellar hemisphere (Fig S2), and not in the adjacent host hemicerebellum, nor in vehicle treated grafts, suggesting that the action of BDNF was local within the denervated tissue.

### BDNF-induced olivocerebellar reinnervation requires PSA-NCAM

To identify *how* BDNF induced sufficient climbing fibre outgrowth over several millimetres to reach and reinnervate Purkinje cells in the grafted hemicerebellum, we examined molecules associated with neural circuit development and plasticity. Polysialic acid (PSA)-NCAM is highly expressed in early development, promoting axonal growth ^**24**^. It is also involved in climbing fibre maturation and post-lesion cerebellar plasticity ^**11**,**25**^.

We measured PSA-NCAM during reinnervation in the cerebellar plate and inferior olive (Table 1). In control tissue, PSA-NCAM concentration decreased during maturation, being approximately halved in the cerebellar plate and inferior olive between 21-26DIV (∼P15-20, Table 1a&b). In lesioned/cocultured explants PSA-NCAM was partially maintained by BDNF application so that by 4 days post-lesion PSA-NCAM was 1.5-2 times greater than control tissue (Table 1b: 26DIV C vs BDNF). Moreover, in a lesioned/isolated cerebellar plate PSA-NCAM expression did not increase after BDNF treatment (Table 1a). Thus, increased PSA-NCAM parallels the greater climbing fibre sprouting in BDNF-treated explants.

**TABLE 1.**
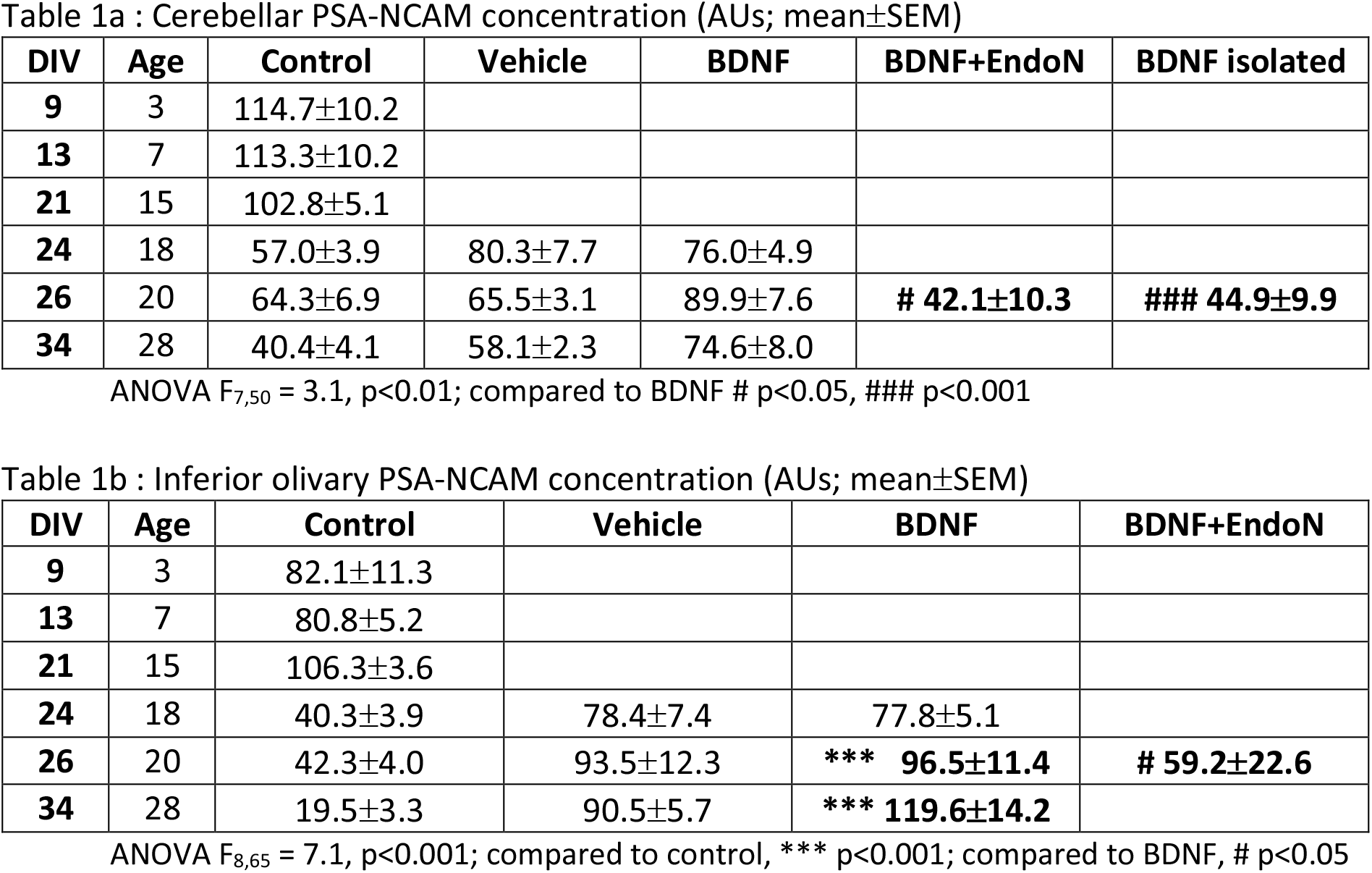
PSA-NCAM concentration in the cerebellum and inferior olive during development and olivocerebellar reinnervation. **A & B:** PSA-NCAM decreases during maturation in the cerebellar plate (A) and inferior olivary region (B). Post lesion, BDNF administration partly counters this decrease, resulting in higher PSA-NCAM tissue concentration, which is significant in the inferior olive (ANOVA F8,65 = 7.1, p<0.001). Additional post-BDNF treatment with the sialase EndoN to lyse PSA from NCAM (BDNF+EndoN) greatly reduces PSA-NCAM when compared to vehicle or BDNF (ANOVA F7,50 = 3.1, p<0.01). Also, post-lesion BDNF does not increase PSA-NCAM in an isolated cerebellar plate, so that there is less PSA-NCAM than in the lesion/coculture/BDNF cerebellum.

To determine whether increased PSA-NCAM is actually involved in olivocerebellar reinnervation, we altered PSA-NCAM concentration in the target (cerebellum) or reinnervating (inferior olive) regions and re-evaluated climbing fibre reinnervation. First, we treated graft hemicerebella with the sialidase EndoN, to remove PSA from NCAM. EndoN reduced PSA-NCAM in lesioned-BDNF-treated hemicerebella (Table 1a) and reduced BDNF-dependant reinnervation by half (Fig 2A). In contrast, overexpression of cerebellar *Sia2*, a sialtransferase that adds PSA to NCAM ^**24**^, induced the same percentage of Purkinje cell reinnervation as did BDNF injection (Fig 2A). However, *Sia2* was not overexpressed in the target Purkinje cells themselves (Fig S3), which suggests that PSA-NCAM acts by providing a growth permissive extracellular environment. Second, we overexpressed *sia2* in the reinnervating inferior olive 24 hours after co-culture (Fig 2B). Olivary *Sia2* overexpression increased climbing fibre-Purkinje cell reinnervation by 50% compared to cerebellar BDNF, but these two treatments did not have additive effects (Fig 2B). In contrast, olivary overexpression of another sialtransferase isoform *sia4*, which is preferentially expressed at mature synapses^**24**^, was less effective (Fig S4). Also, modifying *Sia2* or *Sia4* with a 3’UTR variation to target the mRNA to climbing fibre axon terminals did not further improve reinnervation (Fig S4).

**Figure 2:**
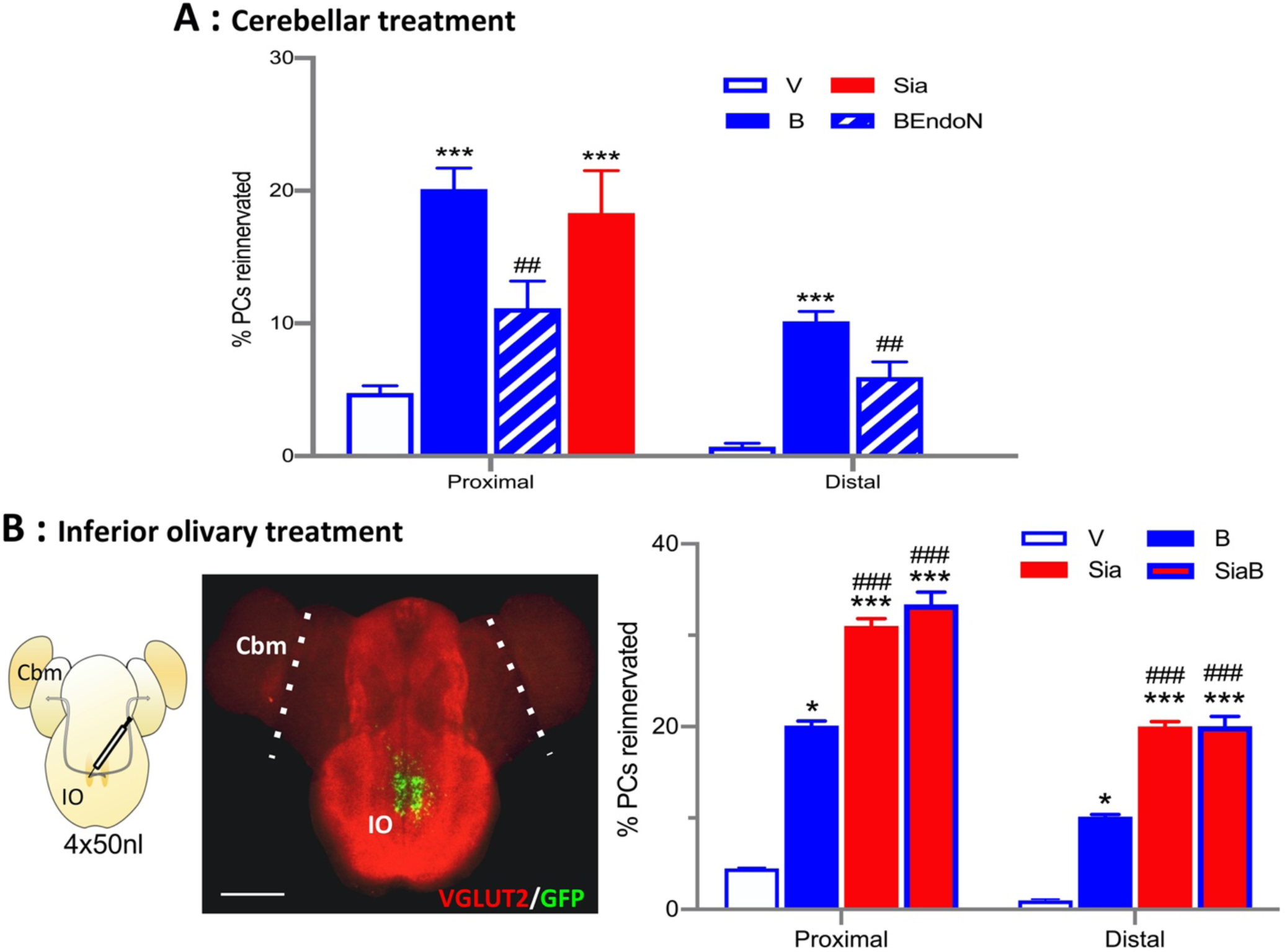
PSA-NCAM induces olivocerebellar reinnervation. **A:** PC reinnervation was analysed at 10 days post-treatment. In the grafted cerebellar plate, reinnervation induced by BDNF (B) is reduced when PSA-NCAM is lysed by the sialase EndoN (B+E; ANOVA, F6,59 = 72.45, p<0.0001) in both proximal and distal zones. Also, overexpression of *sia2* in the cerebellar plate to increase PSA-NCAM, promotes CF-PC reinnervation to a similar level as BDNF (proximal zone sia; F2,27 = 77.11, p<0.0001). Because lentiviral neuronal transfection is slightly variable between explants, we only analysed reinnervation in the proximal zone to avoid variability due to the injection site. Between group post-hoc comparisons BDNF/sia vs. vehicle: ***, p<0.001. Between group post-hoc comparisons BDNF vs. BDNF+EndoN: ##, p<0.01. **B:** Lentivirus injected (4×50 nl) into the explant inferior olive (left panel; IO) is detected by GFP expression (middle panel) or FLAG immunohistochemistry (not shown). The injection remains highly localised to the inferior olivary region. The white dotted line represents the junction between the host and graft hemicerebella (Cbm). Bar = 1mm. CF-PC reinnervation (right panel) in proximal and distal regions of the grafted cerebellar plate, following olivary injection of LV-Sia2 or LV-GFP with (SiaB, B) or without (Sia, V) BDNF applied to the grafted cerebellar plate. In both proximal and distal regions, *sia2* overexpression ± BDNF induced more reinnervation than BDNF or LV-GFP/Vehicle alone (F3,30 = 119.3, p<0.0001); but the 2 treatments were not additive. Between group post-hoc comparisons BDNF/sia/siaB vs. vehicle: *, p<0.05; ***, p<0.001. Between group post-hoc comparisons BDNF vs. sia/siaB: ###, p<0.001.

Taken together these data indicate that BDNF induces climbing fibre reinnervation by increasing PSA-NCAM in the olivocerebellar environment. Increased PSA on the reinnervating axon is highly effective in promoting climbing fibre axon outgrowth and synaptogenesis.

### Paired homeobox transcription factor Pax3 induces olivocerebellar reinnervation

We next looked for the link between BDNF and increased PSA-NCAM. In cerebellar neuronal culture BDNF upregulates *Pax3* ^**26**^, a transcription factor expressed in the immature cerebellum and brainstem ^**23**,**27**^, and which upregulates *Sia2* and *Sia4* ^**28**^. *In silico* searches confirmed the link between BDNF and Pax3 expression, showing connections both through BDNF-TrkB signalling ^**29**,**30**^ and direct protein-protein interactions ^**31**,**32**^ (Fig S5A).

We therefore measured *Pax3* expression in the cerebellar graft and inferior olive 1 - 48 hours after co-culture and cerebellar BDNF treatment. Inferior olivary *Pax3* expression transiently increased 24h post BDNF and was followed by increased Pax3 protein at 48h (Fig 3A, B). There were no changes in Pax3 mRNA or protein in the cerebellum, nor in vehicle-treated controls. These results indicate that increased olivary Pax3 occurs in response to the cerebellar BDNF treatment, at a time when olivary neurons are actively extending axonal branches.

**Figure 3:**
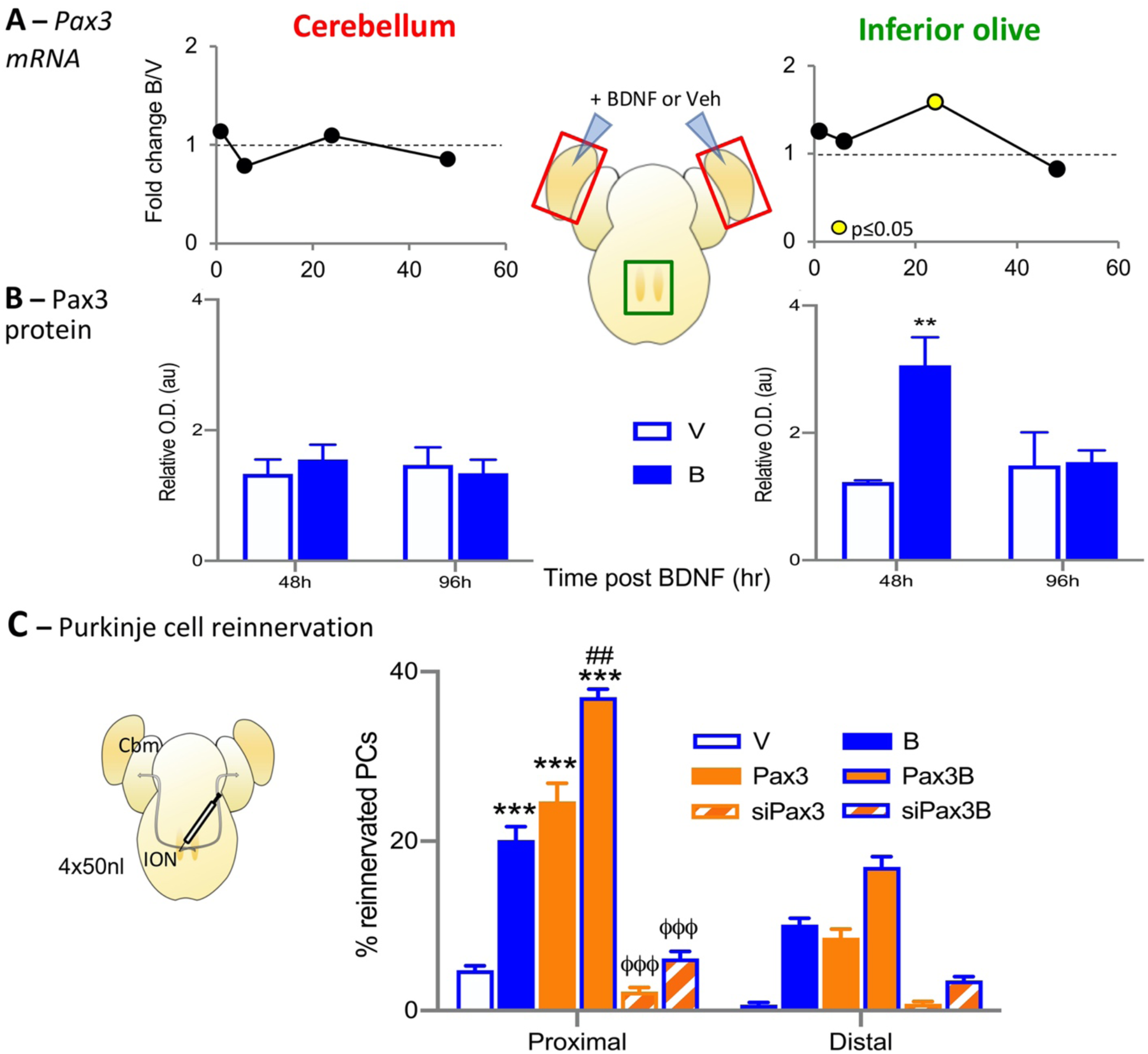
Olivary *Pax3* regulates olivocerebellar reinnervation. **A:** Quantitative PCR of the hemicerebellum or inferior olive 6-48h post lesion and coculture reveals transient *Pax3* upregulation in the inferior olive, but not the cerebellum, 24h after lesion and BDNF treatment (Kruskal-Wallis H8,46 = 16.82, p=0.018; post-hoc MWU: 1h, p=0.55; 6h, p=0.008). **B:** Western blot analysis at 48-96h post coculture also revealed a transient increase in Pax3 protein in the inferior olive after BDNF treatment (ANOVA F7, 40 = 4.170, p=0.0016), but not in the cerebellum. However, increased Pax3 occurred at 48h post BDNF, which follows the rise in RNA (cf A). Between group post-hoc comparisons B vs. V: **, p<0.01; **C:** Modifying inferior olivary *pax3* dramatically alters climbing fibre-Purkinje cell reinnervation (2way-ANOVA, F11,105 = 82.72, p<0.0001). Reinnervation in proximal and distal regions of the grafted cerebellar plate following olivary injection of LV-Pax3 alone (orange bar) or with BDNF (orange with blue border), LV-GFP with (blue bar) or without (white bar with blue border) BDNF. In both proximal and distal regions, *pax3* overexpression induced reinnervation equivalent to BDNF and more than LV-GFP/Vehicle alone (F7,77 = 99.61, p<0.0001). The 2 treatments were additive, so that the effect of LV-Pax3 with BDNF was greater than any single treatment (LV-Pax3 < LV-Pax3 + BDNF, p<0.01). When *pax3* was knocked-down by LV-siPax3 (by 70%, manufacturers data), BDNF-induced reinnervation was decreased (F5,47 = 58.53, p<0.0001; orange diagonal stripes with blue border) to the percentage in vehicle treated explants. Between group post-hoc comparisons BDNF/Pax3/Pax3+BDNF vs. vehicle: ***, p<0.001; Pax3 vs. Pax3+BDNF: ##, p<0.01; siPax3 vs. BDNF/Pax3/Pax3+BDNF: φφφ, p<0.0001. NB: the statistical differences shown in the proximal zone also exist in the distal zone, but for clarity of the figure they have not been added to the graphic.

To test for a role of Pax3 upregulation in olivocerebellar reinnervation, we altered olivary *Pax3* and observed dramatic changes in climbing fibre reinnervation. *Pax3* knockdown effectively abolished BDNF-induced reinnervation (Fig 3C). Conversely, olivary *Pax3* overexpression induced Purkinje cell reinnervation (Fig 3C) with functional climbing fibre synapses (Fig S6). The amount of reinnervation was similar to that induced by BDNF, but *Pax3* was also additive to the effect of BDNF (Fig 3C). Finally, we verified that Pax3-induced effects on climbing fibres resembled those induced by BDNF: 24 hours after olivary *Pax3* overexpression there was extensive sprouting of host climbing fibre horizontal branches (Figure 4A), and cerebellar PSA-NCAM was increased (Fig 4B). The link of these data to olivocerebellar reinnervation is reinforced by the observation of PSA-NCAM on reinnervating axonal growth cones (Figure 4C).

**Figure 4:**
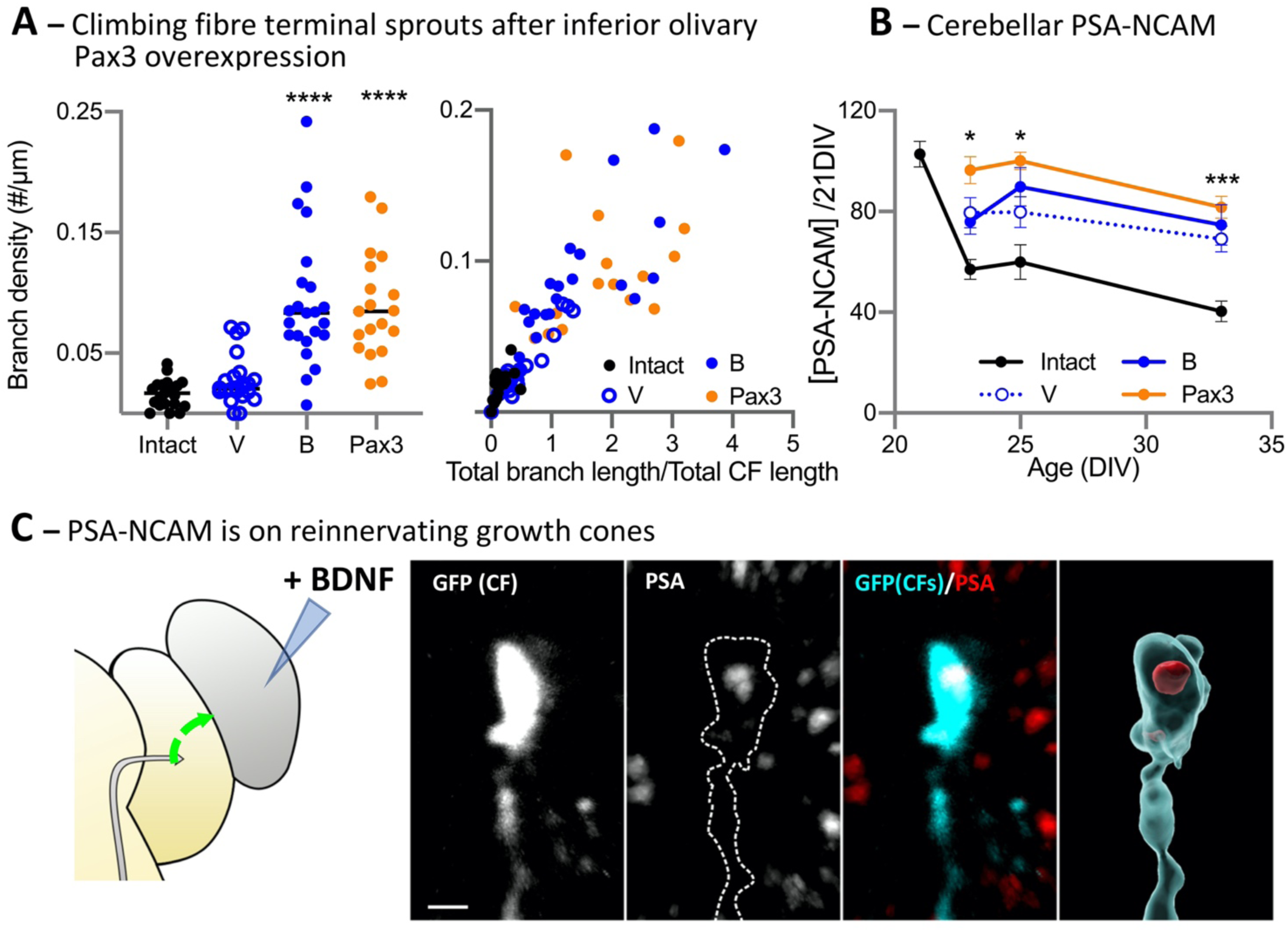
Pax3 increases PSA-NCAM and axonal sprouting. **A:** After lesion and co-culture at 21DIV followed 24h later by olivary LV-Pax3 injection (Pax3), climbing fibre horizontal branches increase within 24h in a manner similar to cerebellar BDNF treatment (B, left panel). There is a greater density of longer branches (Kruskal-Wallis, H4,82 = 52.92, p<0.0001) in BDNF (B) and Pax3 (Pax3) treated groups compared to vehicle (V; Dunn’s post-hoc comparisons, intact vs. BDNF, intact vs. Pax3: both p<0.0001; right panel includes data from Fig 1C for comparison). **B:** As a potential mechanism underlying Pax3-induced reinnervation, we observed increased cerebellar PSA-NCAM from 4 days after lesion and olivary LV-Pax3 transfection when compared to intact or lesion vehicle/LV-GFP controls (F12, 78 = 5.18, p<0.0001) and this was sustained at 12 days post lesion. **C:** In the host cerebellar plate 2 days after lesion/co-culture and BDNF treatment (left-hand drawing), a GFP-labelled olivary axon growth cone, GFP (CF), expresses PSA-NCAM. The right-hand panel is a 3D construction from the z-stack confirming the PSA-NCAM (red) on the growth cone. Bar = 1 µm. Comparisons to intact tissue: *, p<0.05; ***, p<0.001; ****p<0.0001

To verify that Pax3 is a necessary component of BDNF-induced neural circuit repair, we examined its role in repair during development, which occurs spontaneously in the olivocerebellar path ^**7**,**20**,**21**,**33**^ through action of endogenous BDNF ^**34**^. After lesion and co-culture in immature explants equivalent to post-natal day 3 (P3, 9 DIV) to induce spontaneous climbing fibre-Purkinje cell reinnervation (Fig S1), *Pax3* was rapidly upregulated in the inferior olive by 6h (Fig 5A). After ten days, there was robust spontaneous climbing fibre-Purkinje cell reinnervation to over 50% of Purkinje cells in the grafted cerebellar plate (Fig 4B); this result is similar to that seen *in vivo* ^**7**,**20**,**21**^. Moreover, this reinnervation was significantly reduced by either EndoN treatment of the grafted cerebellar plate to reduce PSA-NCAM, or *Pax3* knockdown in the host olivary neurons (Fig 5B). Finally, *in silico* analyses showed that Pax3 target genes are significantly enriched (p<0.0001) in the Gene Ontology functions axon extension (GO:0048675), axon regeneration (GO:0031103) and synapse (GO:0051963; Fig S5B) supporting its role in BDNF-induced neural circuit repair.

**Figure 5:**
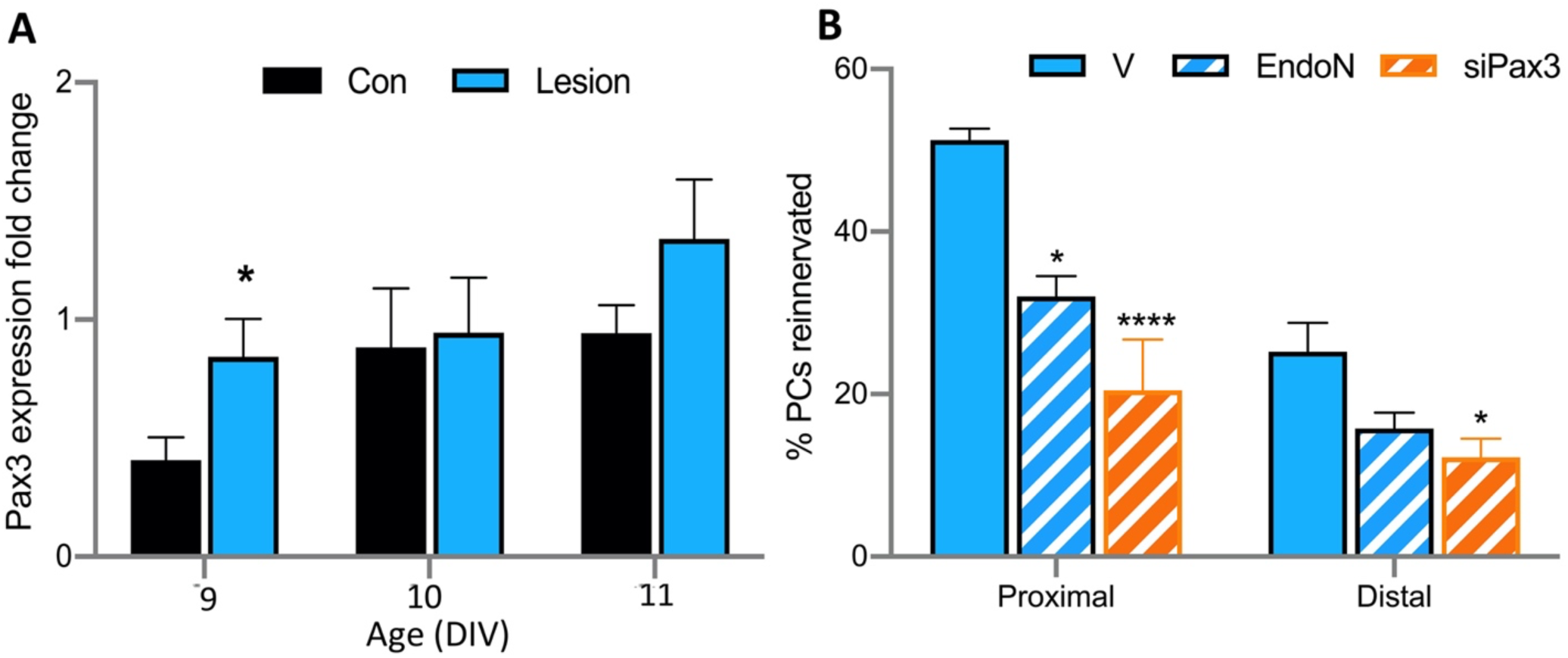
Pax3 is involved in developmental climbing fibre-Purkinje cell reinnervation. **A:** Post lesion and co-culture at 9DIV (∼P3), *Pax3* expression increases very rapidly in the inferior olive (Two-way ANOVA, F2,13 = 4.67, p=0.03), so that at 6h post-lesion there is a 2-fold greater *Pax3* expression compared to controls. This level then remains stable during the period of reinnervation. N=5 olivary regions for each group. **B:** Explants were lesioned and co-cultured at 9DIV (∼P3). The cerebellar plate was then treated with vehicle (V), or EndoN to lyse PSA from NCAM (EndoN), or the inferior olive was injected with LVsiPax3 (siPax3), to knock-down Pax3 expression. The amount of Purkinje cell reinnervation was significantly reduced by both treatments (ANOVA, EndoN: F5,44 = 14.02, p<0.0001). N=cerebellar plates; V = 10, EndoN = 9, siPax3 = 6. Comparisons to vehicle-treated: ** = p<0.01; *** = p<0.0001

These data show for the first time that Pax3 can induce axonal growth and synaptogenesis in the mammalian nervous system and is a necessary component downstream of BDNF.

## DISCUSSION

In this study we used our *ex vivo* model of the olivocerebellar path to understand *how* BDNF promotes post-lesion reorganisation of remaining intact neural circuits to accurately reinnervate denervated targets. We show that BDNF-induced axon collateral growth can be reproduced by presynaptic neuronal expression of the transcription factor Pax3. Pax3 rapidly induces olivary axonal sprouting and subsequent collateral elongation in parallel with increased PSA-NCAM. We also show that both Pax3 and PSA-NCAM are *necessary* for BDNF-induced reinnervation to take place, and that Pax3 stimulates olivary axonal growth by increasing the permissiveness of the axon terminal environment with PSA-NCAM.

### Pax3 induces olivocerebellar axonal outgrowth to reinnervate Purkinje cells

A key discovery of this study is the capacity for Pax3 to induce axonal outgrowth and neural circuit repair. Increased expression of *Pax3* in the inferior olive induced post-lesion Purkinje cell reinnervation by functional climbing fibres, to the same level as that induced by BDNF. Accordingly, *Pax3* knockdown in the inferior olive effectively abolished climbing fibre-Purkinje cell reinnervation normally induced by BDNF application. This is the first demonstration that Pax3 can induce axonal growth and neosynaptogenesis in the mammalian central nervous system.

Pax3 is a paired-box transcription factor that is critically involved in the early development of the nervous system, regulating the proliferation of neural precursors in the neural tube, neural crest and Schwann cell precursors in peripheral nerves ^**35**^. There are no prior reports of links between Pax3 and axonal outgrowth, only its re-expression in damaged Schwann cells after sciatic nerve lesion ^**36**^ that suppresses re-myelination ^**26**,**37**^. Our hypothesis that Pax3 is involved in axon outgrowth is reinforced by two findings.

First, Pax3-induced climbing fibre sprouts and subsequent arbors are morphologically identical to those developed during *in vivo* transcommissural olivocerebellar reinnervation ^**7**,**15**,**21**,**33**^. We show that the source of these collaterals are the horizontal branches of climbing fibre arbors (Fig 1). Thin horizontal climbing fibre terminal branches have been previously described ^**38**,**39**^, although their function is unknown as they do not form synaptic terminals ^**38**^. However, their importance is illustrated by their presence on the newly-formed reinnervating arbors ^**33**,**40**^. The extensive branching of these horizontal branches, induced by either BDNF application or Pax3 expression (Figs 1C and 4B), is entirely consistent with the 15-fold increase in terminal arbors supported by reinnervating transcommissural olivocerebellar collaterals observed *in vivo* in the immature system (up to 89 collaterals per climbing fibre; ^**33**^). Although climbing fibre terminals can readily sprout into the unmyelinated cerebellar molecular layer to reinnervate adjacent denervated Purkinje cells ^**40**,**41**^, BDNF is required for long distance axon collateral extension and climbing fibre-Purkinje cell reinnervation (endogenous BDNF in neonatal animals ^**34**^; exogenous BDNF in the mature system ^**7**,**15**,**16**^). Here we demonstrate that Pax3 is an essential component of this process.

Second, *in silico* analyses revealed that a large number of genes with Gene Ontology functions of axon extension, regeneration, and synaptogenesis, contain Pax3 binding sequences in their promoter regions; and many of these are expressed in the olivocerebellar system (Figure S5B). Our data also provide a mechanism: olivary Pax3 increased PSA-NCAM in the *reinnervating inferior olive*, consistent with Pax3’s upregulation of the sialtransferases *sia2* and *sia4* ^**28**^ inducing a local growth-permissive PSA-NCAM microenvironment on the reinnervating axon growth cones.

### Pax3 allows reinnervating axons to follow directional cues

In addition to stimulating olivocerebellar axon outgrowth through increasing PSA-NCAM, a second key observation is that Pax3-induced reinnervation is target specific: i.e. onto the appropriate cell (Purkinje neuron) and subcellular compartment (proximal dendrites). However, these two phenomena, increased PSA-NCAM and target specificity, are not necessarily concordant.

PSA-NCAM is a canonical component of neurodevelopment and its plasticity, and our data confirm this. Young lesioned explants have high levels of PSA-NCAM (Table 1A) associated with high spontaneous Purkinje cell reinnervation (Fig 5B). Similarly, BDNF application to more mature lesioned-cocultured explants increases PSA-NCAM in the inferior olive in 2 days (Table 1b), when reinnervating climbing fibre collaterals are actively growing ^**21**^, and increased PSA-NCAM, particularly in the reinnervating inferior olive, induces extensive climbing fibre-Purkinje cell reinnervation (Fig 2). Moreover, reinnervation is reduced if PSA-NCAM is removed (by EndoN), and BDNF does not increase PSA-NCAM in the absence of reinnervation, i.e. in an isolated hemicerebellar anlage (Table 1a). However, studies in other brain systems reveal that general increase of PSA-NCAM, for example at a lesion, does indeed promote axonal sprouting, but that reinnervating axons are not of the appropriate phenotype, i.e. are inappropriately targeted ^**42**,**43**^.

In contrast, in our study, the main effect of PSA-NCAM is through its presence on the reinnervating olivocerebellar axons, compared to within the cerebellum. This is illustrated by greater BDNF-induced upregulation of PSA-NCAM in the inferior olive (Table 1B), and greater climbing fibre-Purkinje cell reinnervation when PSA-NCAM is increased in the inferior olive (cf Fig 2A vs B). Although PSA-NCAM is considered a permissive environment that allows axon outgrowth ^**11**^, there is increasing evidence that its presence on the *growth cone* is essential for directing axon extension, both during development ^**44**^ and reinnervation ^**45**,**46**^. Given that *in vivo* climbing fibre-Purkinje cell reinnervation displays appropriate topographic organization ^**20**,**21**,**33**^, we suggest that increasing Pax3 expression in afferent neurons, to increase PSA-NCAM on reinnervating axon terminals, may provide more effective (i.e. appropriately targeted) neural circuit repair without the adverse effects induced by BDNF administration or by generalized increases in PSA-NCAM throughout the neural pathway.

### Pax3 provides a repair program rather than recapitulating development

It is often assumed that neural repair mechanisms recapitulate those of development. However, axonal injury alters the expression of thousands of genes, some of which are required *only* for regeneration/repair and have no known role in development ^**47**^. Moreover, simply promoting mechanisms of neural development is rarely sufficient to induce axon growth and repair in the adult mammalian brain ^**48**^. The identification of Pax3 in our system as an integral part of the BDNF repair program is entirely consistent with these previous findings.

First, Pax3 does not appear to have a critical role during normal olivocerebellar axonal maturation. Pax3 is involved in early neural precursor development ^**35**^, and is strongly expressed in the embryonic mouse hindbrain from E8.5, where it regulates regional developmental genes ^**23**^. *Pax3* expression ceases by E12-13 ^**49**^, a time when cerebellar macroneurons are differentiating, but their migration and axonal growth have barely begun ^**50**^. Second, we show that Pax3 is necessary for repair at different maturation stages. Not only does BDNF upregulate *Pax3* expression in neuronal culture ^**26**^, but we observe Pax3 upregulation in the inferior olive both by BDNF treatment after lesion/coculture at 21DIV *and* during reinnervation after lesion/coculture at 9DIV, which occurs spontaneously through the action of endogenous BDNF ^**34**^. Moreover, spontaneous reinnervation after a developmental lesion (coculture at 9DIV; ∼P3), of which collateral outgrowth from remaining axons is a major component ^**5–7**^, is blocked by knock-down of inferior olivary *Pax3* (Fig 5B). Also, inferior olivary *Pax3* overexpression in the maturing system induces rapid sprouting of climbing fibre horizontal branches within 24h (Fig 4B), which is consistent with rapid spontaneous sprouting and reinnervation post-lesion *in vivo* ^**21**,**51**^.

Thus, our data showing Pax3 upregulation by lesion/BDNF, and its necessity for extensive axon collateral growth, is consistent with a program that correctly guides axons to accurately repair a neural circuit.

## Conclusion

BDNF induces axon collateral outgrowth and targeted neosynaptogenesis in olivocerebellar reinnervation, both in vivo and ex vivo and in developmental and maturing systems ^**7**,**15**,**16**,**19**^. We show that this reinnervation depends on the transcription factor Pax3 and its downstream target, PSA-NCAM. Our data indicate that BDNF induces Pax3 expression which triggers PSA-NCAM synthesis; when present on reinnervating axon terminals, PSA-NCAM promotes target-directed axon growth that contributes to neural circuit repair. This is a novel role for Pax3, but its ability to activate phenomena of directed axon outgrowth, in both developmental and mature systems, opens the possibility of its application to a range of clinical challenges.

## ACKNOWLEDGEMENTS

This project was funded by the Institut pour la Recherche sur la Moelle épinière et l’Encéphale, and the Fondation pour la Recherche Médicale (N° DEQ20081213987). We thank Prof Allan Tobin for all his valuable advice on the manuscript.

## Author contributions

Conceptualization: RMS and JSJ; Methodology: JSJ, HXA, RMS; Investigation: JSJ, HXA, IK, JB, CG, CD, AML, RMS; Data analysis: JSJ, HXA, IK, MD, AML, RMS; Supervision: RMS; Funding and resources: JM, RMS; Writing – original draft: JSJ, HXA, CD, AML RMS; Writing – review & editing: JSJ, HXA, AML, RMS.

## Statement of Interests

The authors declare no competing interests.

## STAR METHODS

### RESOURCE AVAILABILITY

#### Lead Contact

Further information and requests for resources and reagents should be directed to and will be fulfilled by the Lead Contact, Rachel M Sherrard

#### Materials Availability

This study did not generate new unique reagents.

#### Data and Code Availability

This study did not generate new code.

## EXPERIMENTAL MODEL AND SUBJECT DETAILS

### Animals

Timed-pregnant RjOrl:Swiss mice were purchased from Janvier (France) and housed under standard laboratory conditions with 12 hours light/dark cycle and free access to food and water. Age and number of mouse embryos used for each experiment are detailed in the figure legends. Sex of embryos used was not tested. All housing and procedures were approved by the Comité National d’Ethique pour les Sciences de la Vie et de la Santé (N° 1492-02) and performed in accordance with the guidelines established by the European Communities Council Directive (2010/63/EU).

## METHOD DETAILS

### Organotypic cultures

Mouse hindbrain explant cultures were performed at E14, as described previously ^**19**,**22**^. Briefly, pregnant mice were anaesthetised with isoflurane and euthanised by cervical dislocation. Embryo hindbrains (including the cerebellar anlage and the inferior olive nucleus), were isolated and cultured as described. In each litter, explants were semi-randomly assigned to different experimental groups, so that each litter contributed to several groups and each group contained explants from different litters.

### Cerebellar denervation and explant injection

Cerebellar plates were removed from their brainstem at 9 or 21 DIV (equivalent to P3, when olivocerebellar reinnervation is spontaneous, or P15, when reinnervation requires BDNF ^**7**^) and apposed (graft) to intact (host) cerebellar tissue for co-culture. In this configuration, reinnervating olivary axons must grow through white matter and pass neurons of the deep cerebellar nuclei, which is similar to in vivo post-pedunculotomy repair ^**33**^.

#### 21 DIV co-culture

Twenty-four hours after co-culture at 21 DIV, some cerebellar plates were treated with 1µl (4µM) recombinant human brain derived neurotrophic factor (hBDNF; Alomone, 0.1% BSA in H2O). Some co-culture explants also received other treatments: (1) 350 IU of neuraminidase EndoN (AbCys, France) injected onto the graft (denervated) hemicerebellum on +1 (22 DIV) and +7 (28 DIV) days to digest PSA from PSA-NCAM; (2) lentiviral mediated gene transfer to induce over-expression of Sia2, Sia4, Pax3, or GFP in either the graft cerebellar plate (1µl) or inferior olive (4×50nl), injected +1 day after co-culture (22 DIV; see method for virus production below); (3) inferior olivary injection (4×50nl) of lentivirus containing pooled siPax3/hPax3/Pax3i/GFP (LVsiPax3; Gentaur, France) to knock-down Pax3 expression, injected +1 day after co-culture (22 DIV).

#### 9 DIV co-culture

Grafted hemicerebella of some explants were treated, as above, with: (1) cerebellar Endo-N at +1 (10 DIV) and +7 (16 DIV) days; or (2) inferior olivary injection of LVsiPax3 at +1 day (10DIV).

### Recombinant lentiviral vector production

Recombinant plasmid vectors encoding for *sia2, sia2-3’UTR, sia4, sia4-3’UTR, pax3 or enhanced green fluorescent protein (GFP)* under the control of the PGK promoter were used to prepare stocks of lentiviral particles as previously described ^**53**^. All constructs were FLAG-tagged to allow for the identification of transduced cells using a FLAG antibody. Briefly, HEK 293T cells were transiently cotransfected with the p8.91 encapsidation plasmid ^**54**^, the pHCMV-G (Vesicular Stomatitis Virus pseudotype) envelope plasmid and the pFlap recombinant vectors containing the transgene. The supernatants were collected 48 hours after transfection, treated with DNAseI (Roche Diagnostics) and filtered before ultracentrifugation. The viral pellet was then resuspended in PBS, aliquoted and stored at - 80°C until use. The amount of p24 capsid protein was determined by the HIV-1 p24 ELISA antigen assay (Beckman Coulter, Fullerton, CA). Virus from different productions averaged 175 ng/µl of p24 antigen.

### Immunohistochemistry

We used double or triple fluorescent immunolabelling to identify the presence or absence of climbing fibre reinnervation, virally-transfected neurons, and different cell populations in the cerebellum or inferior olivary nucleus (primary antibodies are identified in the Key Resources Table). Fixed whole explants or 10 µm cryosections were incubated with different combinations of primary antibodies, then immunolabelling was visualized with FITC-, Cy5-, or Cy3-conjugated species-specific donkey secondary antibodies (1:200; Jackson ImmunoResearch Laboratories, West Grove, PA) as described ^**19**^. Finally, sections or explants were examined using epifluorescence or confocal microscopy.

### Histological Analysis

The amount of climbing fibre reinnervation was measured 10 days after treatment (with BDNF, EndoN, or LV injection). We quantified the percentage of calbindin-positive Purkinje cells (soma and primary dendrites) colocalised with VGLUT2 per field of view. This quantification was made systematically on z-stacks taken in rows through the cerebellar graft with increasing distance from the host-graft interface. Climbing fibre quantification was made on these z-stacks, hiding each colour channel as necessary to ensure VGLUT2 puncta were not missed. Data from rows 1 and 2 were defined as a proximal zone, and those from rows 3-5 were defined as a distal zone.

### Protein analysis: Western Blot and ELISA

Cerebellar plates or inferior olivary tissue were taken from explants during development from 9 - 35 DIV and at 1, 2, 4, or 12 days post-lesion. Tissue was homogenized in 500 µl of lysis buffer (pH; 120 mM NaCl; 50 mM TRIS, 1 mM EDTA; 25 mM NaF; 1 mM Na3VO4; 0.5% SDS), sonicated and centrifuged (14 000 rpm, 30 min) and the supernatants were analyzed by ELISA or western blot.

PSA-NCAM concentration was measured using an ELISA kit (PSA-NCAM ELISA ABC0027B, Eurobio France) according to manufacturer’s instructions. Samples were run in duplicate. Pax3 protein was measured by western blot; 30 μg of total protein was separated by SDS-PAGE, transferred onto PVDF membranes and probed with anti-Pax3 (1:250; R&D systems) and then anti-β-actin (1:1000; Abcam). Bound antibody was visualised using HRP-conjugated anti-mouse secondary antibody (Amersham Biosciences) and ECL Advance for chemiluminescence (2 minutes exposure time; Amersham Biosciences). Bands were identified by size (Pax3: 53 kDa; βactin: 42 kDa), intensities were measured (ImageJ, Gelplot macro) and normalised against the amount of β-actin in each lane.

### qRT-PCR

RNA was extracted from either the cerebellar plate or the inferior olivary region of co-cultured or control explants 6 - 96h post lesion and co-culture. Tissue from 6 cerebellar plates or inferior olive regions were pooled to obtain each sample. Total RNA was extracted using Trizol (Life Technologies) according to manufacturer’s instructions ^**55**^ and RNA concentration was measured by a NanoDrop 1000 Spectrophotometer (Thermo Scientific, Waltham, MA, USA) before being stored at -80°C.

200ng of total RNA was reverse transcribed in 20µl using a High Capacity cDNA Reverse Transcription Kit (Applied Biosystems). cDNA was amplified on a LightCycler® 480 (Roche Applied Bioscience, USA) in 10 μl reaction volume using SYBR Green I Master Mix (annealing temperature 58 °C, 50 cycles). Housekeeper genes were TUB5 and ARBP and genes of interest were *cfos* and *Pax3*. Primer sequences for each gene are found in the Key Resources Table.

All samples were amplified in triplicate and the mean was used to calculate gene expression in each tissue sample. Raw data were pre-processed with Lightcycler 480 software (Roche Applied Bioscience, USA). Target gene expression was normalised to the harmonic mean of 2 housekeeper genes.

### Electrophysiology

Whole-cell patch-clamp recordings from Purkinje cells in explants were performed as previously described ^**22**^. Patch pipettes were filled with a solution containing: 120 mM Cs-D-Gluconate, 13 mM biocytin, 10 mM HEPES, 10 mM BAPTA, 3 mM TEACl, 2 mM Na2ATP, 2 mM MgATP, 0.2 mM NaGTP, pH 7.3, 290–300 mM mOsm. Explants were continuously perfused with a bath solution containing: 125 mM NaCl, 2.5 mM KCl, 1.25 mM NaH2PO4, 26 mM NaHCO3, 2 mM CaCl2, 1 mM MgCl2, 25 mM glucose, and bubbled with 95% O2 and 5% CO2. Picrotoxin (100 µM) was added to block inhibitory currents. Climbing fibre-EPSCs were elicited by stimulation with a saline-filled glass pipette in the area surrounding the Purkinje cell. We distinguished climbing fibre-EPSCs from currents mediated by parallel fibres (PF-EPSCs) by their all-or-none character and by the demonstration of paired-pulse depression.

Following electrophysiological recordings, explants were fixed in 4% paraformaldehyde (PFA) in PB 0.1 M and processed for immunohistochemistry; the recorded (biocytin-filled) Purkinje cells were visualized after incubation with AF488-conjugated avidin (Invitrogen, Molecular Probes).

## QUANTIFICATION AND STATISTICAL ANALYSES

Statistical analyses were done using GraphPad Prism 9. Data were examined for normality and homogeneity of variance and, where necessary, transformed. Inter-group comparisons were made using analysis of variance (One- or two-way ANOVA) and Bonferroni’s post hoc tests; or Kruskall Wallis and Dunn’s post hoc tests (if normality was not attained). The number of climbing fibre branches was assessed using the χ^2^ test. All values were stated as mean±SEM and α = 0.05. Statistical information (test and p value) is given in the figure legends.

*In silico* searches were used to identify potential Pax3 function in reinnervation. To analyze Pax3’s protein-protein interaction network, we used the STRING database (Search Tool for the Retrieval of Interacting Genes, http://www.string-db.org/). This database integrates text mining in PubMed, experimental/biochemical evidence, co-expression, and database association to provide an interactive platform in which connections, associations, and interactions between proteins can be assessed. An interaction network chart with a combined score >0.8 was saved and exported, and is found in Figure S5A. To complement this analysis, we combined our results with Gene Ontology (GO) terms (GO:0031103 Axon Regeneration; GO:0051963 synapse; GO:0048675 Axon Extension; GO:0044295 Axonal Growth Cone). The web-based service Pubmed was used to perform text mining, and a homemade perl script extracted all available literature related to the search query for co-citation of Pax3 and the genes from GO list. Finally, to explore the putative transcription factor-target relationships of Pax3, we used R (3.3.0) and the Bioconductor suite (3.3). We applied Biostrings (2.40) and GenomicFeatures (1.24) to determine potential transcription factor binding sequences in a selected list. Transcription factor binding matrices [position weight matrix (PWM)] were obtained from the R package MotifDb (1.14) and compared, using matchPWM (with position at 90%) to target sequences in the promoter site and for 2kb upstream.

## SUPPLEMENTAL FIGURES TITLES

Figure S1: Climbing Fibre reinnervation *in vitro*

Figure S2: BDNF action is restricted to the injected graft hemicerebellum

Figure S3: Cerebellar injection of LV-Sia2 does not transfect Purkinje cells

Figure S4: LV-Sia4, 3’UTR-Sia2 and 3’UTR-Sia4 induce olivocerebellar reinnervation

Figure S5: Biological pathways linking BDNF and Pax3

Figure S6: Pax3-induced olivocerebellar reinnervation forms functional climbing fibre-Purkinje cell synapses.

## Notes

### Competing Interest Statement

The authors have declared no competing interest.

