## Supplementary Material for "Pax3 repairs a neural circuit through a program of directed axon outgrowth"

Figure S1: Climbing Fibre reinnervation *in vitro*

Figure S2: BDNF action is restricted to the injected graft hemicerebellum

Figure S3: Cerebellar injection of LV-Sia2 does not transfect Purkinje cells

Figure S4: LV-Sia4, 3'UTR-Sia2 and 3'UTR-Sia4 induce olivocerebellar reinnervation

Figure S5: Biological pathways linking BDNF and Pax3

Figure S6: Pax3-induced olivocerebellar reinnervation forms functional climbing fibre-Purkinje cell synapses.

**Figure S1: Climbing Fibre reinnervation *in vitro***

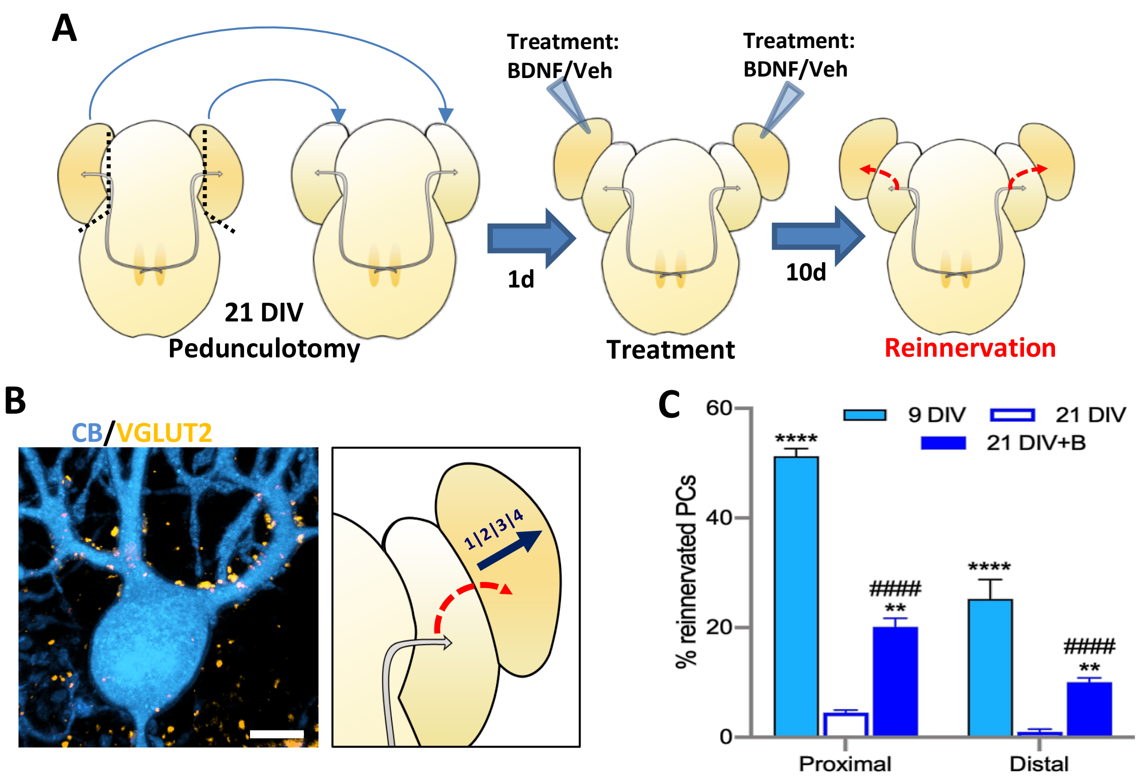


Post-lesion climbing fibre (CF) reinnervation in an *ex vivo* model of the olivocerebellar path mimics *in vivo* repair [S1, S2].

**A:** The E14 mouse hindbrain containing the cerebellum and inferior olivary nucleus is isolated and cultured in "open-book" configuration. Isolated (denervated) cerebellar plates are positioned next to host (intact) cerebella to provide denervated target Purkinje cells (PCs) for reinnervation by the intact olivocerebellar axons (grey arrows). Grafted cerebellar plates can be treated with BDNF or vehicle to induce reinnervation (red dotted arrows).

**B:** Immunostaining for calbindin (CB, PCs) and VGLUT2 (CFs) allows quantification of CF reinnervation in the grafted cerebellar plate following BDNF or lentivirus application to the denervated tissue. CB/VGLUT2 co-labelled PCs are counted on image z-stacks taken in rows with increasing distance from the host (rows 1 to 4). Rows 1 and 2 form the proximal zone; rows 3 and 4 the distal zone. Bar = 10µm.

**C:** Reinnervation of PCs in the grafted cerebellar plate is extensive after lesion at 9 DIV (equivalent to P3) and, following lesion at 21DIV (equivalent to P15), is significantly higher if BDNF is applied (21 DIV+B) compared to vehicle-treated controls (Kruskall-Wallis KW $_{6,69}$ = 63.7, $p<0.0001$). Comparison vs. 21 DIV: ****, $p<0.0001$; **, $p<0.01$ ; comparison vs. 9 DIV lesion, ####, $p<0.0001$.

**Figure S2: BDNF action is restricted to the injected graft hemicerebellum**

To examine the spread of BDNF into the host hemicerebellum, we used *c-fos* expression to detect BDNF action.

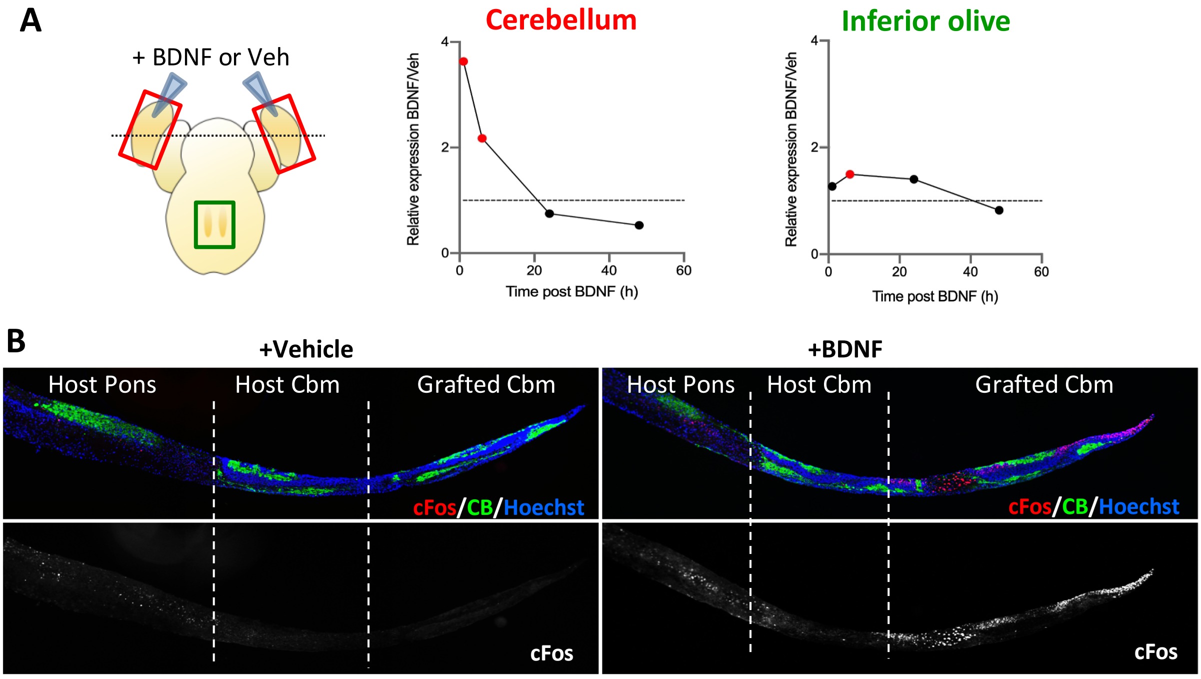


**A:** The diagram (left) shows which tissue was analysed: red squares indicate cerebellar tissue and the green square inferior olive tissue taken for qPCR. The black dotted line shows the orientation of frontal sections of the fixed explant in B. *Cfos* expression fold-change in the cerebellum and inferior olive after BDNF treatment. The horizontal dotted line indicated the control expression in intact host tissue. At 1h post BDNF, there is a 4-fold increase of cerebellar *cfos* (red circle, $p<0.001$), which is still raised at +6h (red circle, $p<0.01$) but has returned to baseline at +24h. In contrast, there is only a small increase in olivary *cfos* 6h after BDNF treatment (red circle, $p<0.01$).

**B:** Immunolabelling shows c-fos (red) after BDNF application to the graft (right panels) but none after vehicle treatment (left panels). In the BDNF-treated explant, c-fos labelling is essentially in the graft cerebellum (right) with minimal expression at the edge of the adjacent host hemicerebellum (centre) and none in the host brainstem (left). This is consistent with the strong cerebellar expression of TrkB receptors [S3, S4] limiting BDNF spread.

The vertical dotted lines indicate the separations between the different areas of the explant.

**Figure S3: Cerebellar injection of LV-Sia2 does not transfect Purkinje cells**

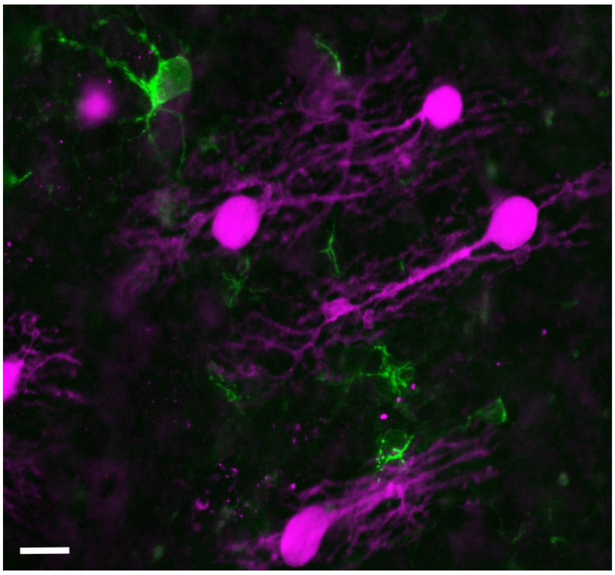

Flattened Z-stack image of the explant hemi-cerebellum after injection of LV-Sia2 counterstained with calbindin to label Purkinje cells (magenta). LV transfection is revealed by Flag expression (green) and this does not co-localize with calbindin. This suggests that any PSA-NCAM synthetized through *Sia2* expression acts through providing a growth permissive cellular environment and does not need to be specifically expressed by target Purkinje cells.
Bar = 20μm.

**Figure S4: LV-Sia4, 3'UTR-Sia2 and 3'UTR-Sia4 induce olivocerebellar reinnervation**

In contrast to LV-Sia2, the effect of LV-Sia4 on CF-PC reinnervation was less pronounced, inducing similar reinnervation to BDNF ($F_{8,65} = 69.9$, $p<0.0001$) and less than LV-Sia2 ($p<0.001$). However, the percentage of PC reinnervation increased when LV-Sia4 and BDNF were combined (Sia4B > Sia4; $p<0.01$) to equal that induced by LV-Sia2 +/- BDNF (Sia, SiaB). Since Sia4 is expressed preferentially in the mature nervous system, specifically in regions of synaptic plasticity [S5, S6], its induction of less reinnervation than Sia2 is consistent with different roles for these two enzymes in the developing/mature nervous system.

Given the large distance between the medullary inferior olive and sprouting cerebellar CF axon terminals, we tested whether targeting the upregulated mRNA to the axon terminal would produce a greater effect. However, inferior olivary injections of LV-Sia2-3'UTR (Sia3') or LV-Sia4-3'UTR (Sia43') in lesioned co-culture explants had only the same effect as the non-3'UTR vectors.

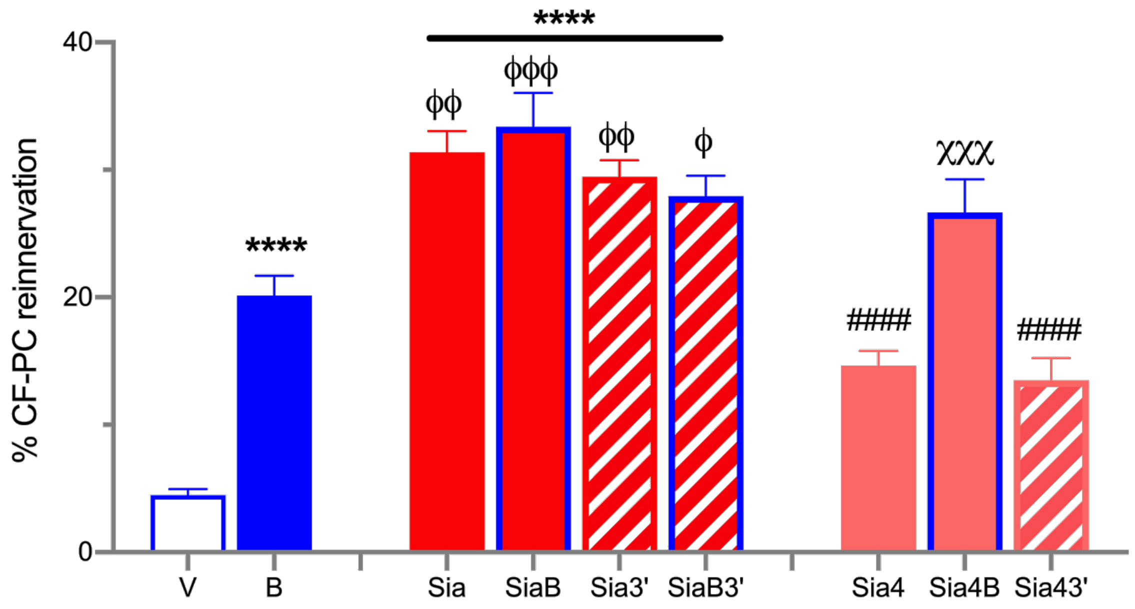


Bars show mean ± SEM

Compared to vehicle **** $p<0.0001$;

Compared to Sia2, ####, $p<0.0001$

Compared to BDNF, ϕ $p<0.05$; ϕϕ $p<0.01$; ϕϕϕ $p<0.001$

Compared to Sia4 and Sia43', χχχ $p<0.001$

**Figure S5: Biological pathways linking BDNF and Pax3**

A potential link between BDNF and PSA-NCAM, which is synthesised through the action of sialtransferases *Sia2* and *Sia4*, is the transcription factor Pax3 [S7]. It is known that in cerebellar neuronal culture BDNF upregulates *Pax3* [S8], which is expressed in the immature cerebellum and brainstem [S9,S10]. Also, BDNF-TrkB signalling induces Klf4 [S11] and Stat3 [S12], both of which bind to response elements in the *Pax3* promoter. Moreover, BDNF has direct protein-protein interactions leading to the Pbx1/Meis2 complex [S13] that directly induces *Pax3* expression [S14]. We used *in silico* searches to confirmed these links.

**A:** Diagram showing BDNF and Pax3 protein-protein interactions from the STRING database with an interaction score of >0.8

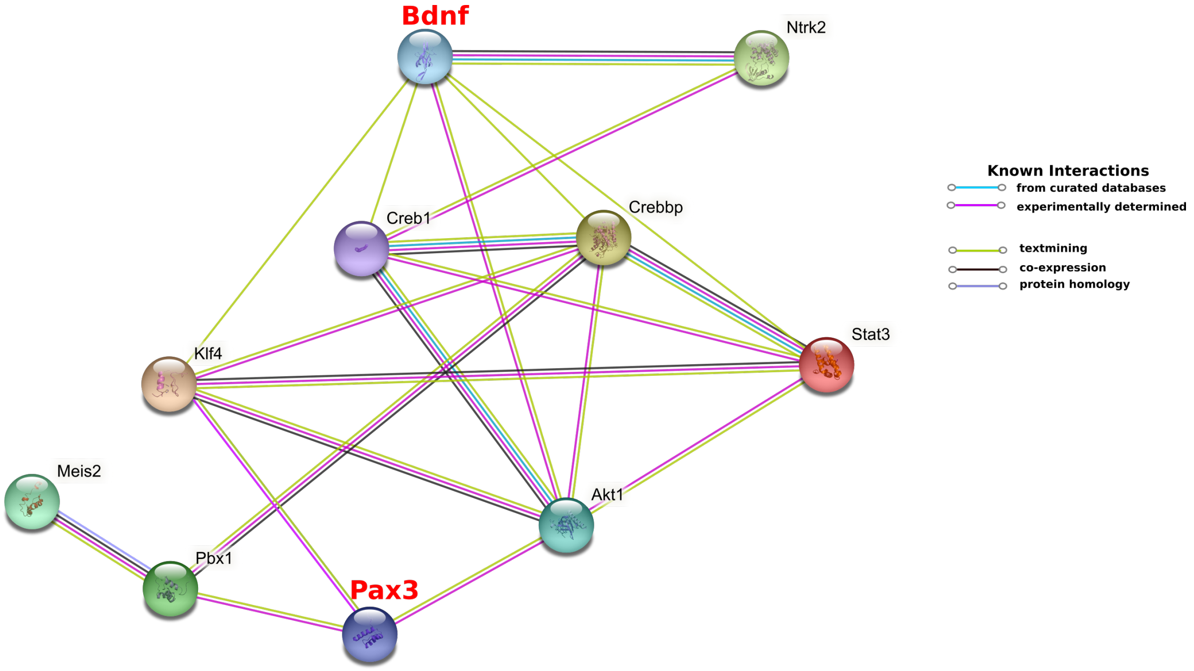


**B:** To clarify that Pax3 was capable of inducing axonal outgrowth and olivocerebellar reinnervation, we searched Gene Ontology functions of interest (axon extension, GO:0048675; axon regeneration, GO:0031103; and synapse, GO:0051963) for genes containing predicted Pax3 binding elements. (see Excel file Jara_Pax3 target genes). The position weight matrix for Pax3 (top left) indicates the probability of specific bases at each position within the response elements. We then used PubMed text mining to identify potential Pax3-targets which are expressed in the olivocerebellar system (in red bold).

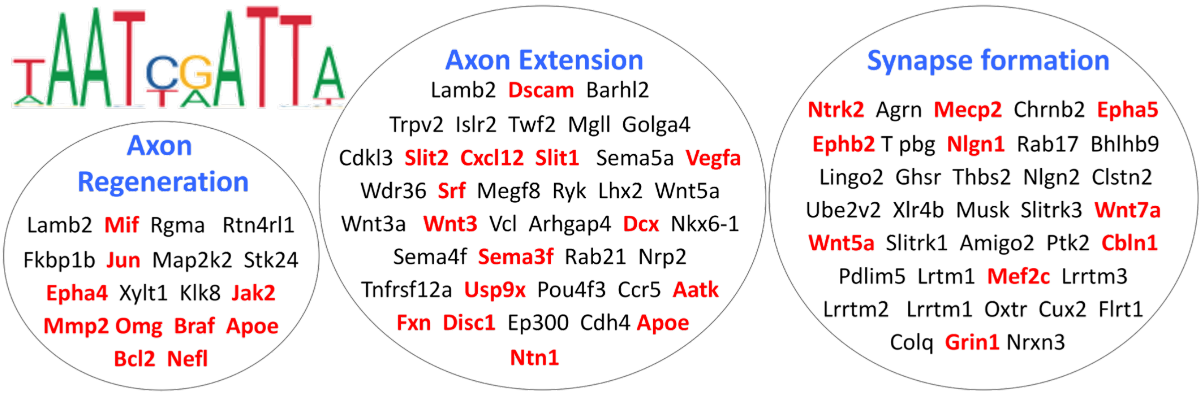

**Figure S6: Pax3-induced olivocerebellar reinnervation forms functional climbing fibre-Purkinje cell synapses.**

Because inducing axonal growth and reinnervation is a new function for Pax3, it was important to confirm that the VGLUT2-labelled terminals contacting cerebellar PCs were in fact functional climbing fibre synapses.

**A:** Whole-cell PC recordings (see Supplementary Methods, below), in the graft cerebellar plates of explants overexpressing olivary *Pax3,* allowed us to find PCs with characteristic CF synaptic currents (CF-EPSCs), showing classic all-or-none activity and paired-pulse depression. Other PCs had no CF-EPSCs.

**B:** Histological analysis revealed that CF-EPSC-positive PCs, identified with biocytin-Alex Fluor488, colocalised with VGLUT2-positive profiles (red; white arrows indicate co-localisation). PCs without recorded CF-EPSCs generally did not have associated VGLUT2 labelling.

Bar = 20 μm

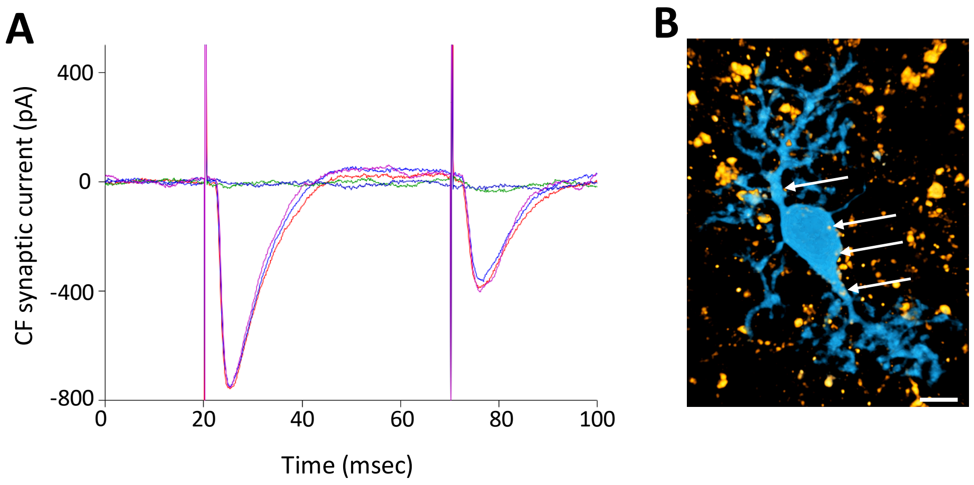
